# Aging alters the functional connectivity of motor theta networks during sensorimotor reactions

**DOI:** 10.1101/2023.07.17.549324

**Authors:** Juliana Yordanova, Michael Falkenstein, Vasil Kolev

## Abstract

**Objective:** Both cognitive and primary motor networks alter with advancing age in humans. The networks activated in response to external environmental stimuli supported by theta oscillations remain less well explored. The present study aimed to characterize the effects of aging on the functional connectivity of response-related theta networks during sensorimotor tasks.

**Methods:** Electroencephalographic signals were recorded in young and middle-to-old age adults during three tasks performed in two modalities, auditory and visual: a simple reaction task, a Go-NoGo task, and a choice-reaction task. Response-related theta oscillations were computed. The phase-locking value (PLV) was used to analyze the spatial synchronization of primary motor and motor control theta networks.

**Results:** Performance was overall preserved in older adults. Independently of the task, aging was associated with reorganized connectivity of the contra-lateral primary motor cortex. In young adults, it was synchronized with motor control regions (intra-hemispheric premotor/frontal and medial frontal). In older adults, it was only synchronized with intra-hemispheric sensorimotor regions.

**Conclusions:** Motor theta networks of older adults manifest a functional decoupling between the response-generating motor cortex and motor control regions, which was not modulated by task variables. The overall preserved performance in older adults suggests that the increased connectivity within the sensorimotor network is associated with an excessive reliance on sensorimotor feedback during movement execution compensating for a deficient cognitive regulation of motor regions during sensorimotor reactions.

**Highlights:**

- The connectivity of motor theta networks is modulated by sensory and cognitive variables in sensorimotor tasks.
- Motor theta oscillations of young adults are synchronized between the primary motor cortex and cognitive control regions.
- In contrast, motor theta networks of older adults are decoupled from motor control regions during sensorimotor reactions.

## 1. Introduction

Motor function declines with increasing age in humans. Major deficits refer primarily to motor slowing, and impairment of fine movements and coordination (Salthouse, 2000; Krampe, 2002). Neuroscience research has since long explored the physiological correlates of these motor alterations with aging (revs. Rossini et al., 2007; Seidler et al., 2010). In addition to peripheral muscular impairments (Clark and Taylor, 2011; Manini et al., 2013), substantial changes have been established at the central level of motor functioning (Ward and Frackowiak, 2003; Andres-Hanna et al., 2007; Seidler et al., 2010), including neuroanatomical (Raz et al., 2009) and neurotransmission alterations (Volkow et al., 1998; van Dyck et al., 2008). However, it has been demonstrated that aging-related functional alterations can contribute to motor dysfunctions beyond the impairments caused by the degradation of the neural matter (Hoffstaedter et al., 2015).

The functional status of the motor system is currently being explored within the concept of distributed motor networks collectively subserving motor functions at the behavioral level (Tomasi and Volkow, 2012; Ash and Rapp, 2014; Antonenko and Flöel, 2014). According to one major distinction, there are two fundamental motor networks: a motor execution network and a cognitive motor control network. The motor execution network involves the primary motor (M1) and sensorimotor cortices and sub-cortical structures - the basal ganglia and the cerebellum (Rizzolatti et al., 1998). The cognitive motor control network involves extended cortical regions in the dorso-lateral and medial frontal and parietal lobes – the premotor cortex, the supplementary motor area (SMA), and the anterior and posterior cingulate cortex (ACC, PCC) (Xiong et al., 1999; Hanakawa et al., 2003, 2005; Hanakawa, 2011; Heckner et al., 2021). Another major distinction identifies motor neural networks based on their primary engagement with externally or internally guided movements (Wu et al., 2004; Wu and Hallett, 2005; Hanakawa, 2011). Generally, medial cortical structures are included in the system controlling intentional and internally motivated movements (Deiber et al., 1999; Hoffstaedter et al., 2014), whereas lateral frontal and parietal regions are more involved in movements relying on guidance by external environmental information, thus mediating the interaction of the organisms with the external environment (Haggard, 2008).

In this perspective, it has been demonstrated that older adults consistently manifest decreased within-network and increased between-network functional connectivity leading to lower network segregation, modularity, and efficiency (rev. Deery et al., 2023; Damoiseaux, 2017). These reports align with existing models according to which aging is characterized by a network reorganization, loss of specificity of the functional systems (dedifferentiation), and inclusion of distributed rather than focused networks in relation to specific functions (Cabeza, 2002; Cabeza et al., 2002; Reuter-Lorenz, 2002; Li et al., 2001; Koen and Rugg, 2019). However, these age differences are reliable for higher-order functional networks, whereas findings for primary sensory and motor networks are variable (Deery et al., 2023).

It is to be noted that many important results about motor networks in aging are derived from observations of resting-state networks (e.g., Stumme et al., 2020; Tian et al., 2018; Samogin et al., 2022). Importantly, the extent to which patterns of functional connectivity are highly correlated between rest and motor/cognitive task states depends on age. Specifically, increasing age is accompanied by differences between the involvement of resting-state and motor-task networks pointing to reduced network flexibility (Monteiro et al., 2019), reduced activation selectivity or differentiation (Chan et al., 2017), and exacerbation of the reduced segmentation of age-related network during motor tasks (Hughes et al., 2020). Thus, studies of networks during motor performance may have a particular independent contribution to understanding aging-related functional networks.

Motor performance in old individuals has been associated with decreases in both the regional activity and effective connectivity within the core motor network engaging the primary motor cortex (Michely et al., 2018). During bimanual motor control, older adults have manifested changes in inter- and intra-hemispheric connectivity between prefrontal and premotor areas (Loehrer et al., 2021), enhanced inter-hemispheric connectivity of the electroencephalographic (EEG) beta-band network (Shih et al., 2021), and stronger premotor-motor connections of EEG beta networks focused in the left hemisphere (Babaeeghazvini et al., 2019). While age-related observations during motor tasks indeed appear diverse, the involvement of beta and alpha networks is detected more consistently. This possibly reflects the established role of alpha and beta event-related synchronization/desynchronization in movement regulation (e.g., Neuper et al., 2006), corresponding to the nature of the bimanual synchronization motor tasks employed.

On the other hand, when processes of external sensory guidance or motor planning, initiation, and sensorimotor integration are considered, networks from slower frequency bands appear to be functionally reactive. Rosjat et al. (2018, 2021) have identified age differences for a low-frequency (delta/theta) network characterized by reduced variability of the subnetworks, emergence of additional enhanced inter-hemispheric connections, and different activations of prefrontal and premotor areas (see Frolov et al., 2020, Pitsik et al., 2022 for similar results). These low-frequency networks appear of special interest also because response-related potentials (RRPs) generated after external sensory stimulation are composed of slower frequency components from delta/theta frequency bands in both young and old individuals (Popovych et al., 2016; Liu et al., 2017; Yordanova et al., 2004, 2020). Moreover, the connectivity between primary sensorimotor/motor and medial frontal regions during movements is supported by synchronized networks from delta and theta frequency bands (Urbano et al., 1996, 1998; Luu and Tucker, 2001; Duprez et al., 2020; Kolev et al., 2023; Yordanova et al., 2023).

Despite the vast range of investigations as outlined above, studies have insufficiently targeted the aging-related patterns of connectivity that characterize RRPs generated in response to external sensory stimulation in sensorimotor tasks. Also, slower frequency components playing a major role in movement generation, integration and cognitive regulation (Caplan et al., 2003; Cruikshank et al., 2012; Duprez et al., 2020) are less frequently explored. Therefore, the main goals of the present study were (1) to characterize the effects of aging on the functional connectivity of theta oscillations generated during response-related potentials in sensorimotor tasks, (2) to explore both motor executive and motor control networks as represented by the connectivity of primary sensorimotor and medial frontal regions, and (3) to evaluate the effects of sensory and cognitive task variables on aging-related motor theta connectivity.

## 2. Methods

### 2.1. Subjects

A total of 27 subjects were studied. They were divided into two age groups: young (*n* = 14, 5f, mean age 22.5 years, SE = ±1.5), and older (*n* = 13, 6f, mean age 58.3 years, SE = ±2.1). All subjects were healthy, without a history of neurologic, psychiatric, chronic somatic, or hearing problems. They were under no medication during the experimental sessions, with normal or corrected-to-normal vision. Similar to the young adults, the older subjects were involved in social and working activities. The study received approval from the Research Ethics Committee at the Leibniz Institute for Working Environment and Human Factors, Dortmund, Germany. Prior to engaging with the study, all participants gave informed consent in line with the Declaration of Helsinki.

### 2.2. Tasks

Three tasks with different complexity were used: a Simple Reaction Task (SRT), a Go-NoGo task, and a four-choice reaction task (CRT). Four stimulus types represented by the letters A, E, I, and O were delivered randomly with an equal probability of 25%. In the SRT, each stimulus had to be responded to with the middle finger of the right hand, irrespective of its type. In the CRT, the letters A, E, I, and O had to be responded to with the left middle, left index, right index, and right middle fingers, respectively. In the Go-NoGo task, the letters E and O were presented with equal probability, and only the letter O had to be responded to with the middle finger of the right hand, whereas the response to the letter E had to be inhibited. In the present study, all stimulus types were analyzed in the SRT and only O stimulus-response trials were analyzed in the CRT and Go-NoGo conditions. Responses in the three tasks were therefore given with the middle finger of the right hand. The response was a finger flexion. Response force was measured by sensometric tensors while subjects produced the flexion.

Each task was performed in two modalities, auditory and visual. Auditory stimuli (duration 300 ms, intensity 67 dB SPL) were delivered via headphones binaurally, with similar envelopes of the sound pressure waves formed for all stimuli. Visual stimuli with the same duration were shown in the middle of a monitor placed 1.5 m in front of the subject’s face. Inter-stimulus intervals varied randomly between 1440 and 2160 ms (mean 1800 ms). If the response was slower than 700 ms after stimulus onset, a feedback tone was delivered at 700 ms. This tone had to be avoided by the subjects by responding fast enough. The order of tasks was SRT (60 trials for each modality), CRT (60 O-trials for each modality), and Go-NoGo (60 O-trials for each modality). Blocks of auditory and visual series were counterbalanced across subjects.

### 2.3. Data recording and processing

EEG was recorded from 64 channels with Cz as reference, with frequency limits of 0.1–70 Hz, and a sampling rate of 250 Hz. Electrooculographic (EOG) recordings were used for the registration of vertical and horizontal eye movements. Electromyogram (EMG) was recorded to control for muscular artifacts. The mechanogram from the right middle finger was recorded to provide for analysis of motion dynamics and characteristics used in RRP analysis. EEG traces were visually inspected for gross EOG and EMG artifacts. Contaminated trials were discarded along with EEG records exceeding ±100 µV. Slight horizontal and vertical eye movements preserved in the accepted trials were corrected by means of a linear regression method for EOG correction (Gratton et al., 1983). The number of trials accepted for the final analysis for each subject and modality was between 50-55 for SRT, 46-50 for CRT, and 47-51 for Go-NoGo tasks. RRPs were computed with a trigger corresponding to a threshold level of 5 N in the mechanogram, thus discarding incomplete responses. All RRP analyses were performed after a current source density (CSD) transform of the signals providing for a reference-free evaluation (Perrin et al., 1989). Data processing was performed using Brain Vision Analyzer 2.2.2 (Brain Products GmbH, Gilching, Germany).

### 2.4. Time-frequency decomposition

Time-frequency (TF) analysis of RRPs was performed by means of a continuous wavelet transform (CWT). Details of the mathematical procedure are presented in Yordanova et al. (2020). To achieve a reliable analysis of low-frequency components in the time-frequency domain and avoid possible edge effects, 4096 ms-long epochs were used, with the moment of response execution (5 N) being in the center of the analysis epoch. A baseline of 600-800 ms before the response was used. TF decomposition was performed on CSD-transformed single-sweep RRPs. The wavelet family was characterized by a ratio of *f*_*0*_/*σf* = 4, where *f*_*0*_ is the central frequency and *σf* is the width of the Gaussian shape in the frequency domain. The choice of the ratio *f*_*0*_/*σf* was oriented to the expected slower TF components. The analysis was performed in the frequency range 0.5–16 Hz with central frequencies at 0.4 Hz intervals. Basing on previous observations (Yordanova et al., 2020, 2023), the relevant theta TF component was measured with central frequency *f*_*0*_ = 5.5 Hz.

### 2.5. Spatial synchronization

To quantify spatial synchronization, the phase-locking value (PLV) was used (Cohen, 2014a; Keil et al., 2022). As a measure of spatial synchronization, PLVs were computed for the theta TF scale at each time-point *t* and trial *j* according to the equation:

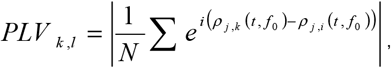

where *N* is the number of single sweeps, *k* and *l* are indices for the pair of electrodes to be compared, and ρ is the instantaneous phase of the signal. *PLV*_*k,l*_ results in real values between one (constant phase difference) and zero (random phase difference). PLV computation followed the approach described in Yordanova et al. (2017). For PLV analysis 35 electrodes were used (F3, Fz, F4, FC5, FC3, FC1, FCz, FC2, FC4, FC6, T7, C5, C3, C1, Cz, C2, C4, C6, T8, CP5, CP3, CP1, CPz, CP2, CP4, CP6, P3, Pz, P4, PO5, POz, PO6, O1, Oz, and O2). PLV was computed for each pair of electrodes, resulting in a total of 595 pairs for each subject, modality, and task (SRT, CRT, Go-NoGo).

For statistical evaluation, only pairs guided by the C3 electrode (n=34; C3-PLV), C4 electrode (n = 34; C4-PLV), and FCz electrode (n = 34; FCz-PLV) were used. Depending on the statistical design (see below), particular pairs were selected from the set of 34 pairs guided by each of these electrodes. The computation of topography maps was done to reflect the topographic distribution of the strength of synchronization of each of these electrodes with the remaining electrodes. In this way, it was possible (1) to analyze separately the connections of the primary motor and medial frontal cortex with specific regions, and (2) to identify regions with strong and weak connections and perform statistical comparisons of the topographic distributions.

### 2.6. Statistical analyses

Theta PLV was measured as the mean magnitude value within -24 to +24 ms around the maximum identified in a time window of -100/150 ms around the moment of response execution. For analysis of *motor execution networks*, response-related theta PLV was analyzed for 20 electrode pairs connecting bilateral frontal, fronto-central, centro-parietal and parietal locations (F3/4, FC3/4, C5/C6, CP3/4, PO5/6) with either C3 or C4 electrode (C3-PLV, C4-PLV). For statistical evaluation a repeated measures ANOVA design was applied with a between-subjects variable Age (young vs. older), and within-subjects variables Contra/Ipsi-lateral hemisphere (CI, C3-guided vs. C4-guided), Task (SRT vs. Go-NoGo vs. CRT), Modality (auditory vs. visual), Region (5 levels corresponding to electrodes at frontal, fronto-central, central, centro-parietal, and parieto-occipital regions), and Laterality (Left vs. Right hemisphere). An additional ANOVA design was performed for 34 pairs capturing with a finer spatial precision the synchronization between oscillations at C3 and all other electrodes, and at C4 and other all electrodes (Age x Contra/Ipsi-lateral hemisphere x Task x Modality x Pair (34 pairs)).

For analysis of *motor control networks*, response-related theta PLV was analyzed for pairs linking the midline fronto-central FCz electrode with bi-lateral electrodes at frontal, fronto-central, central, and centro-parietal, and parieto-occipital cortical areas contra-lateral and ipsi-lateral to the responding hand (F3/4, FC3/4, C5/C6, CP3/4, PO5/6). The same repeated-measures ANOVA design as used for C3-PLV was applied, with the CI variable being excluded. For a more precise assessment of FCz-PLV topography, an additional ANOVA design was applied where a total of 34 pairs reflecting the connections of FCz with all electrodes formed a Pair variable.

## 3. Results

### 3.1. Performance

Statistical results of RT analysis are presented in Table 1(a). As expected, responses were faster in the SRT (232 ± 5.1 ms) and progressively slowed down in the Go-NoGo (301 ± 5.9 ms) and CRT tasks (489 ± 6.8 ms) (Task, *p* < 0.0001). RT slowing with task complexity was more pronounced in older than young adults, as reflected by the significant interaction Task x Age (*p* = 0.002) – Fig. 1. Accordingly, the difference between the two age groups was only significant in the CRT (Age, F(1/25) = 19. 5, *p* < 0.001, *η*^2^ = 0.433). Interestingly, RT was shorter for the auditory than visual SRT (218 ± 7.0 vs. 245 ± 7.3 ms), did not depend on the modality in the Go-NoGo condition (300 ± 8.2 vs. 305 ± 8.6 ms), and was longer for the auditory than visual CRT (495 ± 9.4 vs. 482 ± 9.8 ms) (Task x Modality, *p* < 0.001; Task x Modality x Age, ns), indicating a more robust task complexity effect for the auditory modality in the two age groups. Error rate was negligible in the SRT and Go-NoGo tasks, and did not differ between the age groups for right-hand responses in the CRT.

**Table 1.**
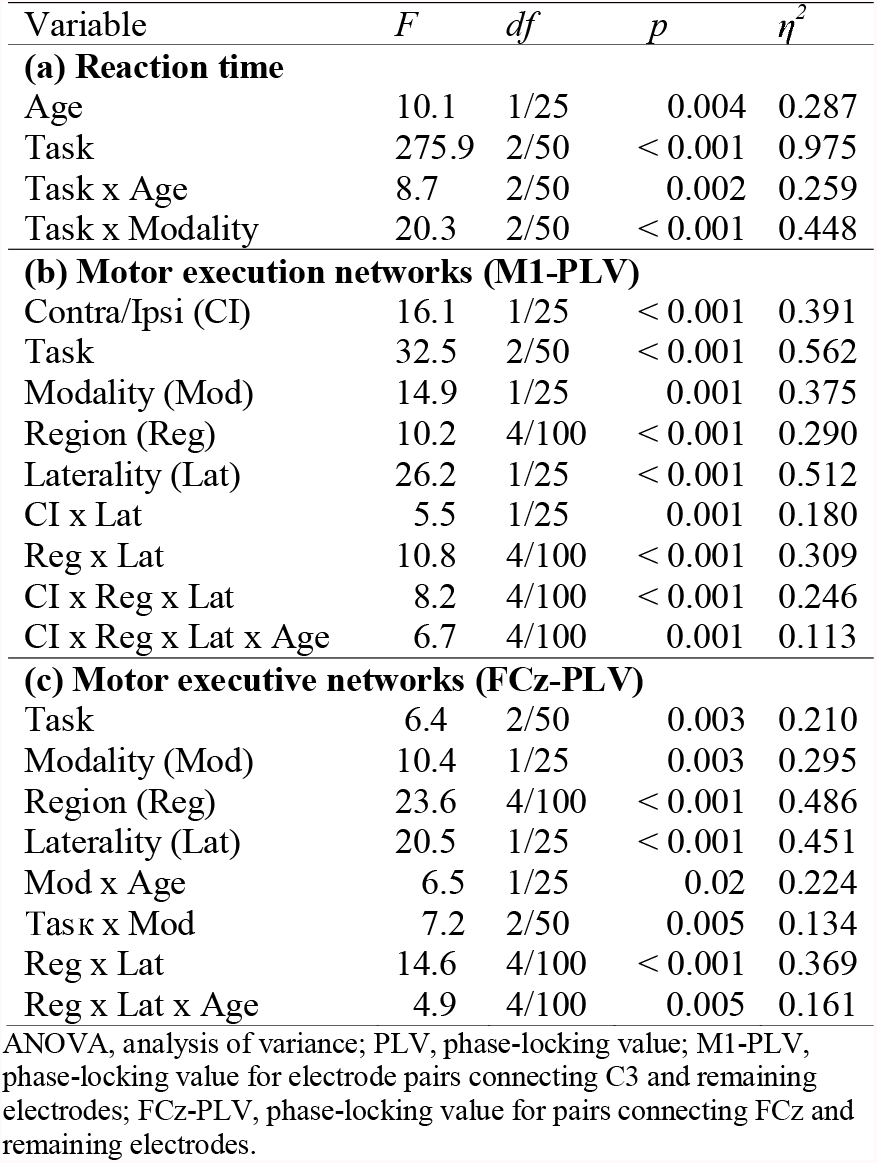
Statistical results of ANOVAs. Only significant main and interactive effects are demonstrated.

**Figure 1.**
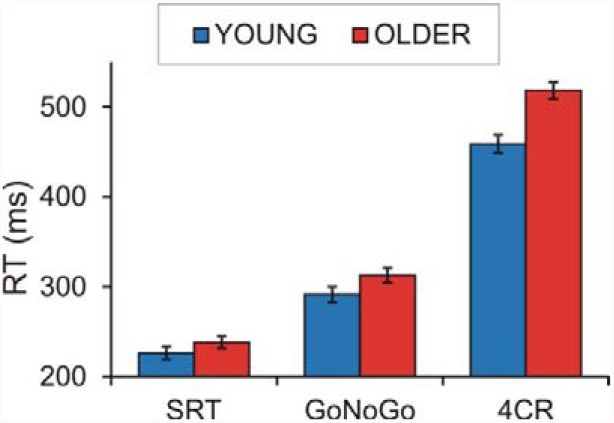
Group mean reaction times (RT) ± SE of young (blue) and older (red) adults in three tasks - Simple Reaction Task (SRT), Go-NoGo task, and four-Choice-Reaction Task (CRT). Auditory and visual modality measures are pooled together.

### 3.2. Motor Execution Networks (M1-PLV)

The following effects illustrated in Fig. 2 were revealed (Table 1(b)). (1) The synchronization of the contra-lateral M1 was significantly stronger than that of the ipsi-lateral M1 (CI, *p* < 0.001). The stronger functional connectivity of the contra-lateral motor cortex was a stable phenomenon, which was not affected by Age, Task, or Modality as indexed by non-significant interactions with CI (*p* > 0.3). (2) The functional connectivity of motor cortical regions was task-dependent, being overall strongest in the Go-NoGo (mean ± SE, 0.598 ± 0.02), moderate in the SRT (0.527 ± 0.02), and weakest in the CRT (0.456 ± 0.026) (Task, *p* < 0.001). The task effect was not modulated by other factors as reflected by non-significant interactions of the Task variable (*p* > 0.2). (3) Interestingly, motor responses in the visual modality were associated with a substantially stronger M1 synchronization as compared with motor responses in the auditory modality (Modality, *p* < 0.0001; 0.553 ± 0.021 vs. 0.501 ± 0.04). (4) During sensorimotor responses the theta connections of M1 were strongest with frontal, fronto-central and centro-parietal areas (Region, *p* < 0.001) in the contra-lateral left hemisphere (Laterality, *p* < 0.001; Region x Laterality, *p* < 0.001; CI x Region x Laterality, *p* < 0.001).

**Figure 2.**
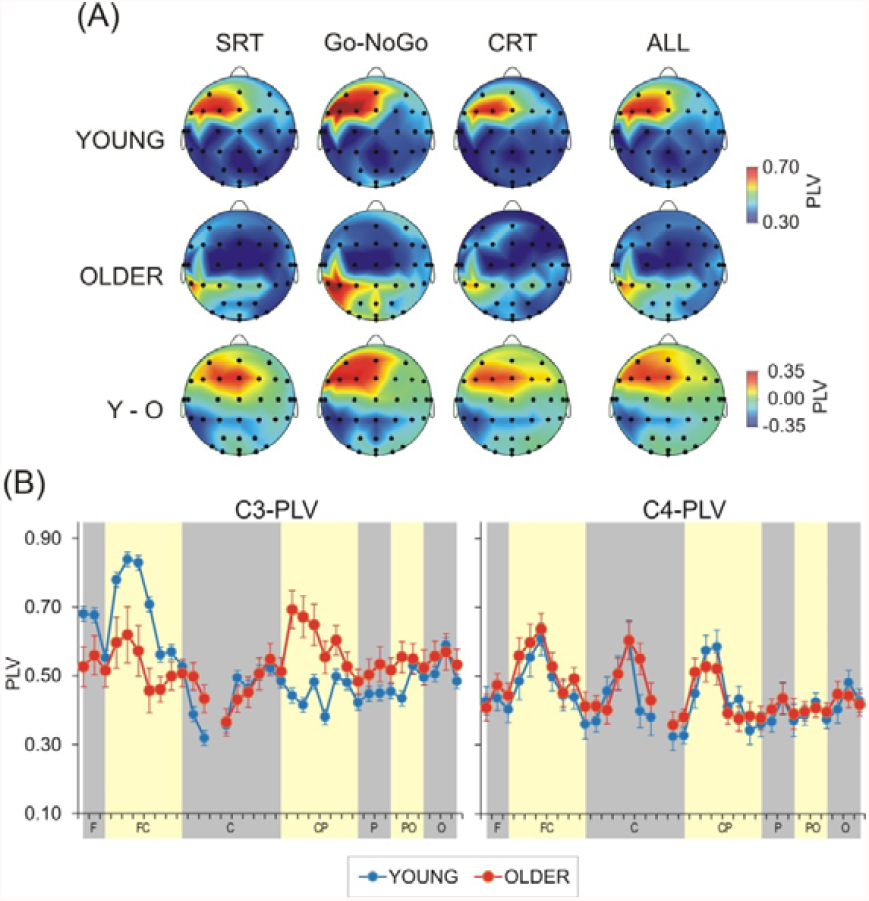
(A) Topography maps of theta C3-PLV at the time of motor response generation computed from response-related potentials (RRP) of young adults, older adults, and young minus older RRP difference in three sensorimotor tasks – a simple reaction task (SRT), a Go-NoGo task, and a four-choice reaction task (CRT). Topography maps reflect the distribution of the strength of synchronization (PLV) between C3 and remaining electrodes. (B) A graphical presentation of C3-PLV (left) and C4-PLV (right) topographic distribution across frontal (F), fronto-central (FC), central (C), centro-parietal (CP), parietal (P), parieto-occipital (PO), and occipital (O) electrodes in young (blue line) and older adults (red line). The arrangement of electrode pairs in each region is from most left to middle to most right. Topography maps and graphical presentation demonstrate that in young adults, motor theta oscillations at C3 are most synchronized with frontal and fronto-central regions of the left hemisphere as well as with medial fronto-central region; in older adults, motor theta oscillations at C3 are most synchronized with sensorimotor motor regions of the left hemisphere. The motor theta synchronization of the ipsi-lateral motor cortex (C4-PLV) is similar in the two age groups. Auditory and visual modality measures are pooled together.

Although no main effect of Age was found (F(1/25) = 0.08, *p* =0.8), aging substantially affected the inter-regional specificity of M1 theta synchronization (CI x Region x Laterality x Age, *p* = 0.001) irrespective of task complexity (interactions with Task, ns). As illustrated in Fig. 2 A, B for C3-PLV, the response generating contra-lateral M1 of young adults was most synchronized with the same-side fronto-central and frontal regions whereas the centro-parietal and parietal regions were not synchronized with either the contra-lateral or ipsi-lateral M1 in young adults (CI x Region x Laterality in young adults, F(4/48) = 8.9, *p* = 0.005, *η*^2^ = 0.428). In contrast, the contra-lateral M1 (C3-PLV) of older adults manifested an increased synchronization with the left centro-parietal regions, whereas no enhancement connectivity with fronto-central and frontal regions was observed (CI x Region x Laterality in older adults, F(4/52) = 5.1, *p* = 0.01, η^2^ = 0.282) – Fig. 2 A,B.

To verify further the effects of aging on the inter-regional connectivity of the response-generating motor cortex, analyses were performed for 34 pairs capturing with a finer spatial precision the synchronization between oscillations at C3 and other regions – Fig. 2B, left. The statistical results of this extended analysis confirmed the main effects of Task (F(2/50) = 26.6, *p* < 0.0001, *η*^2^ = 0.523) and Modality (F(1/25) = 11.4, *p* = 0.002, *η*^2^ = 0.352). More importantly, the Age x Pair interaction was significant (F(33/825) = 4.4, *p* = 0.003, *η*^2^ = 0.183) reflecting a significantly stronger theta synchronization between C3 and fronto-central electrodes FC5, FC3, FC1 and FCz in young than older adults and a significantly stronger theta synchronization between C3 and left centro-parietal sites CP5, CP3, CP1 and CPz in older adults. Testing simple Age effects for pairs linking C3 with separate electrodes confirmed the significant age differences for these electrode pairs (Age, F(1/25) = 7.2–9.8, *p* = 0.01–0.004, *η*^2^ = 0.212–0.293). The ipsi-lateral C4-PLV was equally expressed for fronto-central, central, and centro-parietal pairs (Pair, F(33/825) = 8.7, *p* < 0.001, *η*^2^ = 0.258) and did not depend on age (Age x Pair, *p* > 0.5) – Fig. 2B, right.

### 3.3. Motor Control Networks (FCz-PLV)

Statistical results of ANOVA are presented in Table 1(c). The significant main effect of Task (*p* = 0.003; Task x Age, ns) revealed that the spatial synchronization of FCz-guieded theta oscillations was overall larger for the Go-NoGo task (mean ± SE, 0.558 ± 0.03) and smaller for the CRT (mean ± SE = 0.477 ± 0.02), being moderately expressed for the SRT (mean ± SE = 0.518 ± 0.03). Additionally, the main Modality effect (*p* = 0.003) revealed that FCz-PLV was significantly stronger in the visual than auditory modality (mean ± SE, 0.548 ± 0.03 vs. 0.488 ± 0.02), but only in young adults (Modality x Age, p = 0.03; Modality effect in young subjects, F(1/12) = 14.4, *p* = 0.003, *η*^2^ = 0.523; Modality effect in older subjects, F(1/13) = 0.3, *p* > 0.5). Also, FCz-PLV was larger for the visual modality only in the SRT and Go-NoGo conditions, but not the CRT (Task x Modality, *p* = 0.005).

FCz-guided connections were significantly stronger for the hemisphere contra-lateral to the response (Laterality, *p* < 0.001), being most enhanced at the centro-parietal regions of the left hemisphere (Region, *p* < 0.0001; Region x Laterality, *p* < 0.0001) – Fig. 3A. This distribution was not modulated by processing conditions (interactions with Task and Modality factors, ns) and was observed in each age group (Region x Laterality in young adults, F(4/48) = 12.6, *p* < 0.001, η^2^ = 0.500; in older adults, F(4/52) = 6.4, *p* < 0.001, η^2^ = 0.331) – Fig. 3A. The main effect of Age was not significant. However, in contrast to older adults, young adults manifested a strong synchronization of FCz-guided oscillations not only at contra-lateral centro-parietal, but also at central motor areas (Region x Laterality x Age, *p* = 0.005).

**Figure 3.**
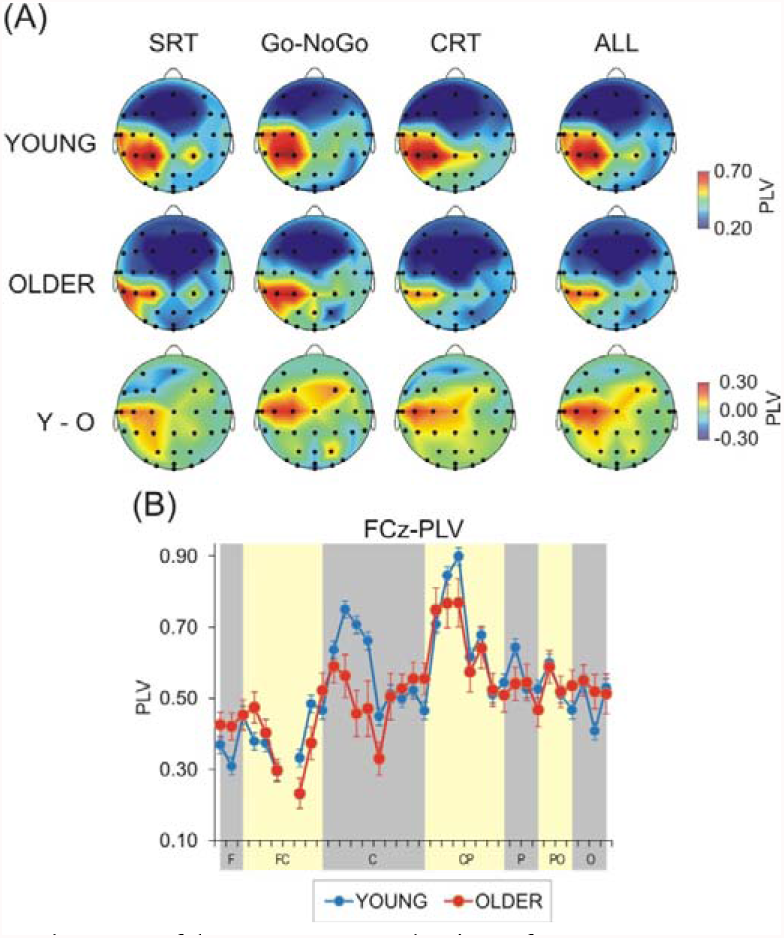
(A) Topography maps of theta FCz-PLV at the time of motor response generation computed from response-related potentials (RRPs) of young adults, older adults, and young minus older RRP difference in three sensorimotor tasks – a simple reaction task (SRT), a Go-NoGo task, and a four-choice reaction task (CRT). Topography maps reflect the distribution of the strength of synchronization (PLV) between FCz and remaining electrodes. (B) A graphical presentation of FCz-PLV topographic distribution across frontal (F), fronto-central (FC), central (C), centro-parietal (CP), parietal (P), parieto-occipital (PO), and occipital (O) electrodes in young (blue line) and older adults (red line). The arrangement of electrode pairs in each region is from most left to middle to most right. Topography maps and graphical presentation demonstrate that in young adults, motor theta oscillations at FCz are more strongly synchronized with the primary motor cortex generating the movement (central left electrodes) as compared to older adults. Auditory and visual modality measures are pooled together.

The refined analysis with 34 pairs confirmed the main effects of Task (F(2/50) = 6.4, *p* < 0.004, η^2^ = 0.205), Modality (F(1/25) = 10.8, *p* = 0.003, *η*^2^ = 0.301), and topography (Pair, F(33/825) = 19.7, *p* < 0.001, *η*^2^ = 0.441). The topography effect again stemmed from a significantly stronger synchronization between theta activity at FCz and centro-parietal areas (CP5, CP3, CP1, CPz). However, the significant Age x Pair interaction (F(33/825) = 2.5, *p* = 0.01, *η*^2^ = 0.100) reflected the enhanced FCz-PLV for C5, C3, and C1 electrodes only in young adults, and a significant reduction of the FCz-guided connectivity in older adults at the contra-lateral primary motor region (Age effect at C5, C3, and C1 electrodes, F(1/25) = 4.7–7.2, *p* = 0.04–0.01, *η*^2^ = 0.160– 0.223; Age effect for the remaining pairs, ns) – Fig. 3B.

## 4. Discussion

The present study was undertaken to characterize the effects of aging on the functional connectivity of motor networks supported by oscillatory theta activity. The effects of aging were analyzed with respect to (1) movements produced in response to external environmental stimuli, and (2) the relationships between motor connectivity and task processing conditions (sensory modality and task demands). As detailed below, the functional connectivity of both motor executive and motor control networks was substantially reorganized in older individuals. These aging alterations persisted across task and modality conditions indicating a stable aging effect on the motor theta networks that was not modulated by functional variables.

In line with a variety of neuroimaging and neurophysiological studies in humans having documented the functional asymmetry during unilateral movements (e.g., Tanji and Mushiake, 1996; Rossini and Dal Forno, 2004; Bhattacharjee et al., 2021; Bundy and Leuthardt, 2019), the contra-lateral motor cortex M1 was significantly more synchronized than the ipsi-lateral motor cortex. The contra-lateral M1 manifested strongest connections with pre-motor and sensorimotor regions in the same hemisphere, whereas the strongest connections of the ipsi-lateral M1 were with premotor, motor and sensorimotor areas of the contra-lateral hemisphere. The functional asymmetry was observed in the two age groups. These observations reveal the spatial characteristics of a dynamic theta network supporting motor response generation in response to external stimuli. Also, they validate the measures used in the present study by substantiating the acknowledged inter-hemispheric asymmetry during unilateral movements.

During movement generation in all sensorimotor tasks, the contra-lateral M1 of young adults was connected with *pre-motor and frontal* areas in the same hemisphere. In contrast, the contra-lateral M1 of older adults was not connected with these frontal regions but it was instead most synchronized with the same-side *sensorimotor areas*. Also, the synchronization between the contra-lateral M1 and medial frontal region was suppressed in older as compared to young adults. However, no difference in connectivity patterns between young and older subjects was observed for the functional connections of the ipsi-lateral M1. These observations reveal that increasing age in humans specifically reorganizes the connectivity patterns of the movement-generating contra-lateral motor cortex.

Notably, the aging-related reorganization of motor theta networks was not modulated by differences in task processing conditions, task complexity and modality. The difference in task complexity was experimentally verified here by the progressive slowing of response speed from the SRT to the Go-NoGo and CRT in the two age groups. In line with previously reported task complexity effects in aging (Kok, 2000; Salthouse, 2000), RT slowing in the group of older adults was only found in the most difficult CRT condition but this did not affect the reorganized spatial patterns in older subjects. In addition, age differences were similar for each modality, auditory and visual. Moreover, although task complexity effect was more robust for the auditory than visual modality in the two age groups, the altered connectivity of the movement-generating M1 in aged subjects was not affected by this interaction. Together, these observations support and extend previous studies (Frolov et al. 2020; Rosjat et al., 2021; Pitsik et al., 2022) by revealing that the detected effects of aging on contra-lateral M1 connections are stable, not being modulated by functional variables and task processing demands.

From a neural network perspective, the present observations indicate an aging-related increase in within-network connectivity (strengthened connections between motor and sensorimotor regions) and decreased between-network connectivity (weak connections between motor and pre-motor, lateral frontal and medial frontal regions). Moreover, no additional connectivity was evident in older adults as previously reported in relation to the over-expressed engagement of the ipsi-lateral (e.g., Mattay et al., 2002; Wang et al., 2017; Rosjat et al., 2018; Ricker et al., 2006) or premotor/frontal/ prefrontal regions (e.g., Roski et al., 2014; Monteiro et al., 2019; Heckner et al., 2021). Thus, the acknowledged age-related dedifferentiation or decline in network fragmentation (Stumme et al., 2020; Samogin et al., 2022; Derry et al., 2023) was not supported by the present observations. To address this discrepancy, it is important to note that dedifferentiation has been consistently associated with motor performance decline whereas performance quality and speed were overall preserved in the sample of old individuals studied here. Moreover, high connectivity within the nodes of specific functional networks, and low connectivity to regions outside this network are associated with preserved functions in aging (Antonenko and Flöel, 2014). It is also important to consider the middle-to-old age (mean 58 years) and the active psychosocial status of the older people involved in the present study. Although the reorganization towards more pronounced functional integration has been detected with age progression from young to middle age, the real expression of such dedifferentiation is evident in the old age (Stumme et al., 2020). Hence, the increase in within-motor network connectivity accompanied by a reduction of between-network connectivity reported here may represent an early-stage functional mechanism protecting performance speed and quality against emerging age-related structural (Antonenko and Flöel, 2014; Grefkes and Fink, 2014; Babaeeghazvini et al., 2019) or neuromodulation (Volkow et al., 1998; van Dyck et al., 2008) impairments.

On the other hand, the age-related changes in the connectivity of motor networks may be essentially associated with the functional specificity of the theta-frequency networks analyzed here. Oscillatory networks in the theta band have been identified as a fundamental neurophysiological substrate of integrative cognitive control (Başar et al., 2001; Cavanagh and Frank, 2014), with motor theta oscillations being critically involved. A variety of reports indicate that synchronized theta activity at the motor cortex subserves integrative functions linked to perception, sensorimotor integration, cognitive control, learning and memory (Caplan et al., 2003; Tomassini et al., 2017; Dufour et al., 2018). According to Ekstrom and Watrous (2014), the coupling between movement-related theta networks and memory-related theta networks serves to integrate bottom-up (sensorimotor driven) and top-down (controlled) processing. Likewise, Ramayya et al. (2021) have provided evidence that theta oscillations coordinate distributed behaviorally relevant neural representations during movement. It has been further posited that the medial frontal regions including the SMA and ACC represent a synchronizing hub of a distributed theta system orchestrating behaviorally relevant contexts in movement regulation and cognitive control (Cohen, 2011, 2014b; Duprez et al., 2020).

In the present study, event-related potentials triggered by the motor response (RRPs) were computed, and spatially phase-locked theta oscillations were subsequently extracted. The applications of this approach has validated the response-locked theta component as a fundamental constituent of RRPs in both young and old people (Popovych et al., 2016; Liu et al., 2017; Yordanova et al., 2004, 2020, 2023; Kolev et al., 2023). Since Popovych et al. (2016) have observed phase-locked delta/theta oscillations for both self-paced and reactive movements, they have suggested that the phase-locking of this component is a ubiquitous movement-related signal which is a prerequisite for the preparation and execution of motor actions. Other RRP studies have further demonstrated that the phase-locked theta RRP component at motor areas contra-lateral to the movement as well as the connectivity of motor theta networks are sensitive to performance accuracy implying a role of synchronized motor theta oscillations in error processing and performance monitoring (Luu and Tucker, 2001; Yordanova et al., 2004, 2023; Kolev et al., 2023). These previous findings based on the specific methodological approach used here confirm that theta oscillations that are inherent to movement generation are also coupled within a distributed theta network for movement cognitive control and integration (Ekstrom and Watrous, 2014; Ramayya et al., 2021; Duprez et al., 2020).

In support, the present results demonstrated that in the two age groups, the connectivity patterns of motor theta networks depended on the sensory and cognitive processing conditions.

Specifically, spatial theta synchronization was most enhanced for the Go-NoGo task and was least pronounced for the CRT. Yet, RTs were most prolonged in the CRT and most speeded in the SRT, implying no associations with task complexity. Further, both the CRT and Go-NoGo tasks required stimulus discrimination, sensorimotor integration and response selection, in contrast to the SRT, but the major distinction in motor theta connectivity between the tasks resulted from the Go-NoGo condition, suggesting that stimulus/response selection may not be a major determinant of the strength of motor theta synchronization. In contrast to both the SRT and CRT, active inhibition (the presence of stop trials) was only required in the Go-NoGo task, pointing to the specific increase of motor cortex connectivity to disinhibited reactions. Previous research has consistently acknowledged the associations between frontal theta oscillations and inhibition processes (Huster et al., 2013; Chmielewski et al., 2016; Dippel et al., 2016, 2017; Pscherer et al., 2019; Adelhöfer and Beste, 2020). The present findings not only confirm the associations between theta oscillations and an inhibitory brain networks but they also provide new evidence for the specific enhancement of motor theta connectivity for disinhibited reactions. In addition, they reveal for the first time an enhanced connectivity of motor theta networks during visuo-motor as compared to audio-motor processing perhaps reflecting the critical role of the visual modality in movement planning, monitoring, and adaptation (e.g., Sarlegna and Mutha, 2015; Archambault et al., 2015; Takakusaki, 2017). Finally, the present results demonstrate that the functional sensitivity of motor theta networks to fundamental determinants of integrative motor regulation and control are not affected by the aging processes before 60 years of age.

In view of these integrative functions of motor theta networks linking perceptual, sensorimotor and cognitive regulations of motor actions, the currently observed aging effects on connectivity may have specific functional implications. The premotor cortex in the left hemisphere has been shown to play a critical role in response selection (Rushworth et al., 1997, 2001, 2003). The suppressed connections between the activated motor cortex and premotor/frontal regions in the left hemisphere of older adults observed here suggest that the selected motor representations may be insufficiently engaged to support the emerging movement in these subjects. Also, an age-dependent decline of the functional connectivity of the SMA has been previously found along with an exclusive decoupling within the ACC (Hoffstaedter et al., 2015). In this perspective, the present results of suppressed synchronization between the contra-lateral M1 and medial frontal regions in older adults imply an aging-related deficiency in providing coordinating functional connections with prefrontal, premotor and parietal cortices as well as subcortical regions involved in movement control and monitoring (Hoffstaedter et al., 2014; Duprez et al., 2020). On this background, it may be further suggested that the increased connectivity within the sensorimotor network in older adults reflects an excessive reliance on sensorimotor feedback during movement execution, which may compensate for a reduced movement regulation, monitoring and control due to a decoupling of the medial frontal region.

One limitation of the present study is the small sample size reducing statistical power and preventing results generalization. Also, it has to be emphasized that the PLV used here measures phase synchrony rather than phase lag. Hence, deep sources may co-synchronize distinct cortical areas that may not be necessarily inter-connected (Cohen, 2014a; Keil et al., 2022). Although in the present study the EEG signals were spatially enhanced by means of CSD, it cannot be excluded that subcortical sources known to be involved in movement generation and regulation (Rizzolatti et al., 1998) have contributed to the observed effects of increased connectivity. Yet, using PLV helps to assert that the decoupling of the primary motor cortex from motor control regions in older adults also involves subcortical circuits. Future research is deserved to isolate the role of subcortical networks for inter-regional cortical connectivity in the aging brain.

## Conclusions

By comparing the connectivity of motor theta networks during reactions to external sensory stimuli in young and older subjects, the present study demonstrates that the inter-regional connections of the movement-generating motor cortex contra-lateral to the response are significantly reorganized with increasing age in humans. The ageing-related reorganization is characterized by an intra-hemispheric increase of connections with sensorimotor regions and a decrease of connections with premotor, frontal and frontal medial regions. Independently of age, the connectivity of motor theta networks is enhanced in visual-motor and inhibitory control conditions. However, the aging-related changes in connectivity patterns of motor theta networks persisted irrespective of different task conditions on the background of overall preserved performance in older subjects. These results (1) demonstrate an age-related decoupling of the motor cortex from cognitive control regions during sensorimotor reactions, and (2) suggest that the increased connectivity within the sensorimotor network in older adults reflects a compensatory excessive reliance on sensorimotor feedback during movement execution, and (3) verify the integrative role of motor theta networks in movement regulation and cognitive control. Thus, new evidence is provided that age progression from middle to old age is already accompanied by a deficient cognitive regulation of motor theta networks during sensorimotor reactions.

## Acknowledgment

Work was supported by the National Research Fund of the Ministry of Education and Science, Sofia, Bulgaria (Project DN13-7/2017).

